# AalpiRNA-18529 regulates vitellogenesis of *Aedes albopictus* via the *Gadd45a*-mediated JNK-dependent nurse cell apoptosis pathway

**DOI:** 10.1101/2024.08.20.608798

**Authors:** Lu Yang, Yonghui Gao, Yulan Chen, Shuyi Ren, Yifan Guo, Peiwen Liu, Khadija Batool, Jianxia Tang, Jinbao Gu

## Abstract

*Aedes albopictus* shows a rapid global expansion and dramatic vectorial capacity for various arboviruses. Mosquitoes display distinct sexual dimorphisms, only adult females consume blood meals to complete ovarian follicle development. Therefore, cyclic reproduction in female mosquitoes serves as a foundation for the transmission of numerous disease-causing pathogens. Aedes have an expansion of the piRNA biogenesis genes, indicated that piRNA may play multiple functional roles in mosquitoes. Although the antiviral function of piRNA pathway in mosquitoes has been extensively studied, the role of piRNAs in mosquito reproduction remain to be further understood. In the present study, we first profiled the characteristics of sex-biased piRNAs in adult *Ae.albopictus*. Then, we identified a female biased piRNA (Aalpi18529) in adult females, that was highly expressed in ovaries at blood feeding-dependent termination, and depended on PIWI5 and ago3 mediated biogenesis. Aalpi18529 overexpression suppressed ovarian development, and reduced fertility and fecundity in adult females post-bloodmeal. Furthermore, we demonstrated that Aalpi18529 can effectively repress its direct target, growth arrest and DNA-damage-inducible protein 45a (GADD45A), and eventually regulates ovarian development via the *Gadd45a*-mediated JNK-dependent nurse cell apoptosis pathway. Our study is the first to report an endogenous piRNA, which trigger silencing of an important protein-coding gene by posttranscriptional regulation in mosquitoes, expanding our current understanding of the important and multiple roles of piRNAs in biological processes in *Ae. albopictus*.

**Author Summary:** Here, we conducted high throughput piRNA sequencing and comprehensive analysis of piRNA sex-based expression profiles in adult females and males of Aedes albopictus. Based on several established universal tools for research, we demonstrate an ovary-enriched endogenous piRNA, Aalpi-18529, is involved in the regulation of the apoptosis of nurse cells during vitellogenesis via the GADD45A/phosphorylated JNK (pJNK) axis and ultimately affects ovarian development. In general, uncovering the biological functions of sex-biased piRNAs in Aedes albopictus will enhance the understanding of piRNA roles in mosquito Sexual dimorphism (SD) and will provide provide more information about the high reproductive capacity of Aedes albopictus, which is essential to find alternative control strategies.

**Classification:** Research Reports

## 1. Introduction

Sexual dimorphism (SD) is prevalent in the insect world and refers to any systematic difference in form between males and females of a species, including external appearances, internal organs and biological functions. Generally, these sexually dimorphic traits are grouped into the following three types [1]: 1. primary sexual characteristics, including anatomical and physiological traits that are directly related to sexual reproduction, such as external and internal genitalia and anisogamy; 2. secondary sexual traits, referring to characteristics that are not directly associated with reproduction but increase the probability of reproductive success, including body size, colouration, ornamentation, weaponry and antenna, etc. [2–4] 3. Ecological sexual dimorphism is a characteristic of insects that occupies different habitats of life between the two sexes, and this list of sex traits is most abundant and is enormously diverse in insects [5].

Mosquitoes (Diptera: Culicidae) are undoubtedly the most important insect vectors for the transmission of parasitic and viral infections to humans and animals and still pose serious public health risks. Mosquito vectors display distinct SDs between males and females. In addition to the visible primary sexual and secondary sexual characteristics, mosquitoes display distinguished, sexually dimorphic behaviour between sexes, and only adult females consume blood meals to complete ovarian follicle development. Therefore, cyclic reproduction in female mosquitoes serves as a foundation for the transmission of numerous disease-causing pathogens. However, the evolution of SD is mostly attributable to selective forces from sexual selection [6,7] and ecological roles [8–10]. There is still rather limited knowledge about the explicit molecular mechanisms of the development and evolution of these sexual traits. For certain sexual characteristic traits, a comprehensive understanding requires a systematic elucidation of the complex sex-related gene regulatory networks. Certainly, high-throughput sequencing (HTS) provides valuable insights into the causal relationship between sexually dimorphic traits and sex-related gene expression during embryonic, juvenile and adult development [1].

With the differential expression profiles of noncoding RNAs (ncRNAs) observed between sexes via HTS [11–14], the regulatory roles of ncRNAs in SD have recently attracted growing attention. For example, miRNA-1-3p functions as a male sex-determining factor (M-factor) in the early embryogenesis of *Bactrocera dorsalis* by suppressing its target gene *tra* [15]; miR-2738 plays a regulatory role in the *Bombyx mori* sex determination cascade by interfering with the sex determination genes *P-element somatic inhibitor (PSI)* and *masculinizer (Masc)* [16]. Moreover, a 5’UTR-overlapping lncRNA can directly regulate the expression level of male-specific *dsx* in the water flea *Daphnia magna* (Phylum: Arthropoda, Class: Branchiopoda) [17]. In *Drosophila melanogaster*, a master sex-determination switch gene, *Sex-lethal (Sxl)*, is regulated by lncRNA by influencing the dose-sensitive establishment promoter [18]. In our previous work, 37 lncRNAs were identified that show significant sex-biased expression profiles in the Asian tiger mosquito *Aedes albopictus* [19].

PIWI-interacting RNAs (piRNAs) are single-stranded, 24–31 nt small RNAs and are defined as a third class of small noncoding RNAs (sncRNAs). The highly conserved primary function of piRNAs is to defend genomic identity by repressing transposable elements (TEs) in both germ cells and somatic tissues across most animal phyla. However, in recent years, increasing evidence has revealed that some nonrepetitive and nontransposon-related piRNAs are largely involved in endogenous mRNA regulation by piRNA-mediated mRNA decay and even by increasing mRNA stabilization through polyadenylation [20,21]. Indeed, mRNA regulation is also regarded as an ancestral function of piRNAs [22,23]. Furthermore, the role of piRNAs in the sex determination cascade was recently described [24]. In silkworm, a female-specific piRNA produced from a piRNA precursor feminizing gene *(Fem)* arranged in tandem in the sex-determining region of the W chromosome repressed the Z chromosome-linked *masculinizer (Masc)* mRNA level by means of piRNA-dependent cleavage. *Masc* encodes a lepidopteran-specific zinc finger protein (ZFP), which is involved in the sex-determination pathway in the silkworm by controlling both masculinization and dosage compensation [24–26]. A similar function of piRNAs is also observed in Caenorhabditis, and an X-linked piRNA, *21ux-1* (21U-RNA on the X chromosome), represses *XO lethal–1* (xol-1) expression and then regulates X chromosome dosage compensation and sex determination [27].

The Asian tiger mosquito *Aedes (Stegomyia) albopictus* (Diptera: Culicidae), a mosquito native to Asia, is rapidly expanding its geographic distribution in many countries, is ranked as one of the world’s 100 most invasive species and is being recognized as having increasing importance in the transmission of vector-borne viruses, Dengue virus (DENV), Chikungunya virus (CHIKV) and Zika virus (ZIKV) [28]. Furthermore, the high reproductive capacity of *Ae. albopictus* is an essential factor that contributes to its successful dramatic global expansion. *Ae. albopictus* has a large genome (1.190-1.275 Gigabase pairs. Gb) [29,30], and a lineage-specific expansion of PIWI family genes was observed in *Ae. albopictus* compared to *Drosophila* [31,32] indicated that PIWI proteins may play distinct functional roles in mosquitoes. Although the antiviral roles of PIWI family genes and piRNAs in mosquitoes have been extensively studied in recent years [33,34], the functional diversification of PIWI family genes and the role of piRNAs in mosquito morphology, physiology, behaviour, etc., remain to be further understood. In this study, the sex-biased expression pattern of piRNAs was explored in *Ae. albopictus*, and whether these sex-biased piRNAs are involved in the regulation of ovarian development and fecundity was further determined. This will not only significantly enhance our understanding of an essential role of the piRNA pathway in mosquito SD as well as the molecular mechanism of female reproductive regulation but also offer potential target genes for genetic-based mosquito population control strategies.

## 2. Materials and Methods

### 2.1 Mosquito rearing and maintenance

Colony of *Ae. albopictus* (Foshan strain) originated in Foshan City, P. R. China and was established in the laboratory in 1981. Briefly, mosquito colonies were maintained in humidified incubators at 27 ± 1 °C with 80 ± 5% relative humidity on a 12-h light:dark (LD) cycle. Larvae were reared in pans (50 × 30 × 15 cm) containing 12 litres of deionized water and fed finely ground fish food mixed 1:1 with yeast powder. Adult mosquitoes were maintained in cages (50 × 30 × 30 cm) and constant exposure to 10% sucrose presented through cotton balls as a carbohydrate source. For egg production, adult females were provided defibrinated sheep blood (Thermo Fisher Scientific® Aust Pty Ltd) by using a Haemotek membrane feeding system (Discovery Workshops, Accrington, UK).

### 2.2 Construction and sequencing of a small RNA library

Both unmated adult female and male mosquitoes were collected two days post emergence and pooled (n=15) (glucose fed, blood starved, three biological replicates per group). Total RNA was isolated using TRIzol® reagent (Thermo Fisher Scientific, Waltham, MA, USA) following the manufacturer’s instructions. The purity and concentration of the total RNA samples were determined using a 2100 Bioanalyzer (Agilent) and ND-2000 instrument (NanoDrop Technologies), respectively. Small RNAs were purified from isolated total RNA using 15% denaturing polyacrylamide gel electrophoresis (urea PAGE), and RNA oligonucleotide adaptors were sequentially ligated to the 3’ and 5’ ends of small RNA by T4 RNA ligase (Takara, Dalian, China). cDNA synthesis was performed using oligo (dT) primers with SuperScript III Reverse Transcriptase (Thermo Fisher Scientific), and then a small RNA library was prepared, followed by 18 cycles of polymerase chain reaction (PCR) amplification. Purified cDNA libraries were used directly for cluster generation and sequenced by an Illumina Genome Analyser (Illumina Inc., San Diego, CA, USA). All sequencing was performed by Gene Denovo Biotechnology Co. Ltd. (Guangzhou, China). Raw sequence reads were submitted to the National Center for Biotechnology Information (NCBI) Gene Expression Omnibus (Accession number: GSE266103).

### 2.3 Data analyses

Adaptor sequences were removed, and low-quality tags were cleaned. Contamination due to adaptor-adaptor ligation was removed using Trimmomatic-0.30 with default settings [35]. Unique reads of 24-30 nt were selected for further analysis and mapped to *Ae. albopictus* Foshan strain Aalbo_primary.1 genome (GenBank assembly accession: GCA_006496715.1) by using Short Oligonucleotide Analysis Package 2 (SOAP2) [36]. Reads mapped to rRNAs, tRNAs, snRNAs, snoRNAs, miRNAs, and exons were excluded; reads containing poly-A/T/C/G nucleotides (minimum of 8 homopolymer repeat nucleotides) were also removed. The rest of the genome that matched with small RNAs was used as piRNA-like small RNAs for further analysis.

### 2.4 Stem‒loop RT‒PCR and qRT‒PCR analyses

Candidate piRNAs were reverse transcribed to cDNA using specific stem‒loop RT primers (Table S3). Quantitation was performed on an ABI 7500 Fast Real-Time PCR system (Applied Biosystems Life Technologies, Foster City, CA, USA) using cDNA-specific forwards primer and a universal reverse primer, as listed in Table S3. U6 small nuclear RNA (U6 snRNA) was used for normalization in all samples. Gene expression levels were analysed by the 2^^(−ΔΔCt)^ method [37]. The specificity of amplification was confirmed by melting curve analysis and by running PCR products on agarose gels (1.5-2%).

For sanger sequencing of the piRNA stem-loop RT-PCR products. Candidate piRNAs were reverse transcribed to cDNA using specific stem‒loop RT primers (Table S3). RT-PCR was performed by applying specific primers and the PCR product was runned on agarose gels (1.5-2%) to identify the length. The PCR product was then purified and recovered using GeneJET PCR Purification Kit (Promega, Madison, WI, USA). Then the purified product was ligated with pLB vector (Tiangen, China) and sent to Sangon Biotech (Shanghai, China) for sequencing.

### 2.5 Oligonucleotides and dsRNA

dsRNA was synthesized in vitro using the T7 RiboMAXTM Express RNAi System (Promega, Madison, WI, USA) according to the manufacturer’s instructions. The Aalpi18529 mimics and the Aalpi18529 inhibitor oligonucleotides, as well as the corresponding negative control, were synthesized by GenePharma (Shanghai, China). 2’-O-methylation modification was carried out in all the bases. All dsRNA and oligonucleotide sequences are listed in Table S3.

### 2.6 PIWI-Protein Family Interference

For knockdown experiments, two-day-old *Ae. albopictus* female adults were infected with *Ago3* (GenBank no. Loc109411297) dsRNA, *Piwi5* (GenBank no. Loc109413120) dsRNA and *Vasa* dsRNA (GenBank no. Loc115267274) via intrathoracic injection. Briefly, adults were anaesthetized on ice, and approximately 1 μL of dsRNA (2 μg/μL) was injected into the thorax under a microscope, as described previously [38,39]. After injection, the adult mosquitoes were immediately transferred to small plastic cups (270 ml, 8 cm diameter at the top), fed a 10% glucose solution through soaked cotton wicks and allowed to recover. Red fluorescent protein (RFP)-derived dsRNA (dsRNA-*rfp*) was injected as a negative control for RNAi assays. Three independent biological replicates were included for each treatment (n = 30 per replicate). The inhibitory effects of *Ago3*, *Piwi5* and *Vasa* were quantified by real-time PCR, whereas the transcript levels of Aalpi-529 were measured using stem‒loop RT‒qPCR assays with a SYBR Green kit (Yeasen, China) according to the manufacturer’s instructions.

### 2.7 Generation of antibodies

Peptides containing amino acid sequences of endogenous PIWI proteins (AGO3: (C+)TSGADSSESDDKQSS, (C+)IIYKRKQRMSENIQF;PIWI5: (C+)DIVRSRPLDSKVVKQ, CANQGGNWRDNYKRAI) were synthesized chemically. The peptides were conjugated to KLH, and complete Freund’s adjuvant was used to elicit an immune response. Polyclonal antibodies against AGO and PIWI5 were produced in rabbits by Convenience Biology Corporation (CBC, Changzhou, China). Both polyclonal antibodies against AGO3 and PIWI5 were purified using an ImmunoPure IgG Purification Kit (Pierce, USA) according to the manufacturer’s protocols, and the titre of antibodies was measured using ELISA (Zoonbio Biotechnology Co., Ltd.; Nanjing, China).

### 2.8 Construction of PIWI protein-expressing plasmids

The full-length AGO3 and PIWI5 ORFs with a V5 epitope fused to the C-terminus were subcolonded into the *NotI-NruI* restriction site of AePUb-RFP [40] to create the plasmids AePUb-AGO3-V5 and AePUb-PIWI5-V5, respectively. All plasmid constructions were confirmed by Sanger sequencing.

### 2.9 RNA immunoprecipitation assay

RNA immunoprecipitation (RIP) was modified using previously published methods [41]. Briefly, C6/36 cells were cotransfected with Aalpi18529 mimics and AGO3 and PIWI5 protein expression plasmids (AePUb-AGO3-V5 and AePUb-PIWI5-V5), respectively, and then lysed in radioimmunoprecipitation assay (RIPA) lysis buffer. Next, the cell lysates were incubated with RIPA buffer with V5-Trap® Magnetic beads (Proteintech, Rosemont, IL, USA) conjugated with V5 antibody with rotation at 4 °C overnight. Furthermore, anti-IgG antibody served as a control. After centrifugation, the beads were washed three times with RIPA wash buffer, rinsed with RIPA wash buffer and resuspended in 500 μL of TRIzol LS (Life Technology, USA). After isolation and purification of the immunoprecipitated RNA, qRT-PCR was adopted to quantify RNA enrichment. Enrichment was normalized to 10% input using 2^−ΔCt,^ and the relative enrichment level to the NC-IgG groups was calculated by 2^−ΔΔCt^.

### 2.10 Immunoprecipitation and Western blotting

First, C6/36 cells were transfected with AePUb-AGO3-V5 and AePUb-PIWI5-V5. Two days after transfection, C6/36 cells were lysed with RIPA buffer, and IP was performed with custom-made antibodies against PIWI5, AGO3 (1:10 dilution) and V5 Tag Monoclonal Antibody (1:10 dilution, Invitrogen, USA). The details of the immunoprecipitation procedure were as previously described [42]. For Western blotting, the IP samples were boiled in 6x protein loading buffer for 10 min at 95 °C and then subjected to SDS‒PAGE on a 12% polyacrylamide gel, followed by electrophoretic transfer to PVDF membranes (0.45 mm, Merck Millipore, MA, USA). Membranes were subsequently blocked with 5% bovine serum albumin (BSA) (Genview, USA) in TBST buffer for 2 h at room temperature and then incubated with primary antibodies as follows: Piwi5 (1:10 dilution), Ago3 (1:10 dilution) and V5 Tag Antibody (1:10 dilution). Anti-β-actin rabbit polyclonal antibody (1: 2500, Abcam, UK) was used as a loading control and for normalization. After incubation overnight at 4 °C, the membranes were incubated with the corresponding secondary antibodies (ABclonal, USA) for 2 h at room temperature. Specific protein bands were detected using a ChemiDoc™ Touch Imaging System (Bio-Rad, USA).

For JNK detection, total protein was extracted using RIPA lysis buffer containing protease inhibitor cocktail and phosphatase inhibitor cocktail (Yeasen, Cat# 20109ES). Total JNK, phosphorylated JNK, and β-actin in various treatment and control samples were detected using an anti-human SAPK/JNK polyclonal antibody (Cell Signaling Technology, Danvers, MA, USA; Cat#9252), an anti-phospho-human-SAPK/JNK (Thr183/Tyr185) rabbit monoclonal antibody (Cell Signaling Technology; Cat#4668), and an anti-β-actin rabbit polyclonal antibody, respectively, by Western blotting. The density of phosphorylated JNK was quantified with the image analysis software ImageJ and normalized to that of β-actin. Differences were statistically evaluated using Student’s *t* test in SPSS 17.0 (RRID:SCR_002865).

### 2.11 Ovarian Dissection and Phenotypic Observation

Forty-eight-hour hours postemergence (PE) female mosquitoes were treated with piRNA mimics/inhibitor or dsRNAs as described above. Injected mosquitoes were allowed to recover for 2 days and then were allowed to commercial defibrinate sheep blood (Solarbio Life Sciences, Beijing, China) supplied through Haemotek membrane feeding systems (Haemotek Ltd., Blackburn, UK). Only freshly blood-fed female mosquitoes with obviously engorged, bright red abdomens were selected for subsequent analysis. At 72 h post blood meal (PBM), the ovaries were dissected by tearing off the soft cuticle between the fifth and sixth abdominal sternites with a fine needle and then pulling off and placing the terminal segments in phosphate-buffered saline [43]. The ovaries were dissected, and two indicators were chosen to evaluate ovarian development: (1) the number of developing follicles and (2) the follicle size per individual.

### 2.12 In situ hybridization and staining

Ovaries of 24 h PBM adult *Ae. albopictus* were dissected into phosphate-buffered saline (PBS) solution on ice. Tissue samples were fixed within 20 min of collection for 30 min in 4% paraformaldehyde. After 3 washes for 5 min each in PBST (PBS + 0.1% Tween-20), the paraformaldehyde was completely removed from the solution. Probes in situ hybridization was carried out as described previously [44] with modification using the gene-specific probes shown above and presented in Table S3. Samples were incubated overnight at 37 °C with a specific biotin probe and SA-Cy3 (1:1) (Gene Pharma, China). Nuclei were stained with DAPI (4′,6-diamidino-2-phenylindole) (Gene Pharma, China). Ovaries were mounted in Antifade Mounting Medium with DAPI (Beyotime Biotechnology, Cat#P0131) and imaged 3– 10 days after mounting.

### 2.13 Image acquisition and composition

The fluorescent testes and ovaries were imaged on an All-in-One Fluorescence microimaging system BZ-X800/BZ-X810 (KEYENCE, Japan). Either single slices or image stacks were taken. Images were deconvolved using the microscope’s built-in software (BZ-X800 Viewer).

### 2.14 Dual-luciferase assay

The binding site of Aalpi18529 in the 3’UTR of *AalGadd45α* (LOC109423337), including the upstream and downstream sequences (with a total length of 264 bp) flanking the core binding site of Aalpi18529, was cloned and inserted into the pmirGLO Dual-Luciferase Expression Vector (Promega, China) to construct the WT-GADD45A plasmid. Concurrently, to construct the MUT-GADD45A plasmid, a mutant in the Aalpi529 seed complementary region of WT-GADD45A was generated. For the luciferase assay, HEK-293 cells were transfected with WT-GADD45A (or MUT-GADD45A) and Aalpi18529 mimics (or NC piRNAs). After transfection for 48 h, luciferase expression was assayed using the Dual-Luciferase (R) Reporter Assay System (Promega) according to the manufacturer’s instructions.

### 2.15 Isolation of Nuclear and Cytoplasmic Fractions

For the assessment of the subcellular localization of Aalpi18529, adult female nuclei and cytoplasm were separated using a Cytoplasmic & Nuclear RNA Purification Kit (Norgen Biotek, Ontario, Canada) following the manufacturer’s instructions, and qRT‒PCR analysis was performed as described above. *Ae. albopictus ribosomal protein S7* gene (*AalrpS7*) or *actin-5C* gene (*β-actin*) were used as indicators of cytoplasm-enriched transcripts, and *Ae. albopictus U6 spliceosomal RNA* (*U6*) was used as an indicator of nuclear-enriched transcripts.

### 2.16 Fecundity and Fertility assays

For each treatment group, a total of eight blood-fed inseminated female mosquitoes were released into a cage. The individual oviposition cups were transferred to each group mosquito cage. The number of eggs produced was counted every 24 h for 7 days, and three replicates were performed. Fecundity was evaluated by calculating (i) mean daily egg laying. (ii) Calculative egg number and (iii) total number of eggs laid per female.

To determine the fertility of each female, the hatched eggs were counted with the aid of a light box and magnifier immediately every 24 h after the eggs were submerged and were monitored for 5 days. The number of eggs hatched at each time point was determined by counting 1st instars. The larvae were picked out by pipette every 24 h, and the number of dead and live larvae at each treatment was counted until there were no new larvae. The hatch rate (HR) was calculated as follows: HR% = dead and living larvae/total eggs × 100%.

### 2.17 mRNA Library Construction and Sequences Analysis

First, Forty-eight-hour hours postemergence (PE) female mosquitoes were treated with piRNA mimics as described above. Injected mosquitoes were allowed to recover for 2 days and then were provided blood meal feeding as described above. Only freshly blood-fed female mosquitoes with obviously engorged, bright red abdomens were selected for subsequent analysis. At 72 h post blood meal (PBM), the ovaries were dissected and total RNA were extracted from dissected ovaries (three biological replicates, n = 30 per replicate). The purity and concentration of the total RNA samples were determined using a 2100 Bioanalyzer (Agilent) and ND-2000 instrument (NanoDrop Technologies), respectively. RNA integrity was checked using a 1% agarose gel. After quality control, a certain amount of RNA sample was denatured and later enriched for mRNA with oligo(dT) attached magnetic beads. The mRNA is then fragmented and the reaction system is prepared to synthesize the first and second strands of cDNA. The double-stranded cDNA fragments are then end-repaired, and a single “A” nucleotide is added to the 3’ end of the passivated fragments. Subsequently, PCR amplification reactions are performed and quality control tests are carried out. Single-stranded PCR products are produced via denaturation. Finally, sequencing was performed to generate RNA libraries. All sequencing was carried out at the Beijing Genomics Institute (BGI) (Shenzhen, China). Raw sequence reads were submitted to the NCBI Gene Expression Omnibus (GEO) under the accession numbers GSE266102.

### 2.18 TUNEL Assay of DNA fragmentation in ovarian tissue

Two days after intrathoracic injection of piRNA mimics or dsRNA, the ovaries were dissected and pretreated following a previously described procedure[45] and then subjected to the TUNEL assay to observe apoptosis by using an apoptosis detection kit (Promega, USA, Cat# G3250). The experimental procedures were performed according to the manufacturer’s instructions with slight modifications described previously [45]. For apoptotic cell death analyses. After staining and washing, the samples were fixed on a glass slide using DAPI-containing mounting Vectashield (Beyotime Biotechnology, China, Cat# P0131). Finally, an ultrahigh resolution laser scanning microscope (KEYENCE, BZ-X800/BZ-X810, Japan) was applied to analyse the samples. Cells stained with TUNEL and DAPI fluorescence were regarded as apoptotic cells. For negative controls, ovaries were incubated with labelling buffer without TdT enzyme.

## Results

### 3.1 Global analysis of small RNA libraries and molecular characterization of predicted piRNA-like sRNAs

To explore the sex-based expression profiles of piRNAs in *Ae. albopictus*, small RNA libraries from 2-day-old emerged Virgin females and male adults were constructed by using Illumina Solexa high-throughput sequencing technology. The six libraries produced a dataset of 91,802,246 raw reads in total: 45,595,366 and 46,206,880 reads from females and males, respectively. After removing 5’-adaptors, trimming 3’-adaptor sequences, reads smaller than 18 nt and filtering out low-quality reads, a total of 76,884,188 clean tags were obtained, representing 38,279,848 and 38,604,340 clean tags in female and male libraries, respectively (Table S1).

Reads corresponding to piRNAs were first identified by excluding unique reads of 24–30 nt that mapped to rRNAs, tRNAs, snRNAs, snoRNAs, and miRNAs and reads that contained poly-A/T/C/G nucleotides. A total of 4,751,477 and 3,720,551 reads were identified in female and male libraries, respectively (Table S1), and then mapped to the aligned to the *Ae. albopictus* genome (AalbF2, assembly: GCA_006496715.1, NCBI) [30]. As a result, 269,624 (77,333 are uniquely mapped to the reference genome, whereas 192,291 tags map to multiple locations in the genome) and 160,539 (53,893 are uniquely mapped to the reference genome, whereas 106,646 tags map to multiple locations in the genome) unique reads in females and males, respectively, were selected as piRNA-like sRNAs (pilRNAs) for further analysis (Fig. S1, Table S1) [46]. Considering that it is difficult to further determine and functionally analyse host genes that pilRNAs originate from multiple mapped genomic locations, we focused on uniquely mapped pilRNAs. The general characteristics of uniquely mapped pilRNAs are shown in Fig. S1, and the majority of these pilRNAs were derived from lncRNAs in both females (48.00%) and males (55.73%) and then derived from intergenic regions (31.79% and 24.63%), mRNAs and finally repetitive elements (Fig. S1).

### 3.2 Identification of sex-biased pilRNAs and Experimental Validation

To identify sex-biased pilRNAs, we compared the expression profiles of 2-day-old sugar-fed virgin adult females with those of males. In total, 6471 uniquely mapped pilRNAs displayed sex-biased expression (*P* ≤ 0.05, |log_2_-fold change| ≥1) (Fig. S1F and Table S2). A total of 4208 and 2263 transcripts showed significantly higher abundance in females and males, respectively. Furthermore, 3274 and 1280 pilRNAs showed female-specific (0 reads in males) and male-specific (0 reads in females) expression patterns (Fig. S1F).

To confirm the sequencing data of mosquito pilRNA prediction and identification, the top 5 female (threshold: |log_2_ fold change| ≥2 and female total reads count ≥50) (for Aalpi00028089, Aalpi00005608, Aalpi00006951, Aalpi00006236, Aalpi00001939, Aalpi00026490 and Aalpi00013664 were thought to be different length pilRNA isoforms, we selected Aalpi00005608 as a representative isoform to detect, Fig. S2) and male-biased expressed (threshold: |log_2_ fold change|≥3 and male total reads count ≥150) pilRNAs were selected for stem-loop RT-PCR and qRT-PCR analysis. As a result, all ten pilRNAs were confirmed by stem-loop RT-PCR followed by conventional Sanger sequencing (Fig. 1A and B), except Aalpi00024145 (eight consecutive base variations were detected at nucleotide positions 17-24, Fig.1B). The sequences of pilRNA candidates were entirely consistent with small RNA sequencing data. Seven of ten pilRNAs displayed obviously sex-biased expression patterns in adults by further qRT-PCR analysis. In particular, Aalpi00029231 and Aalpi00018529 showed more than 40-fold higher expression levels in females than in males.

**Figure 1.**
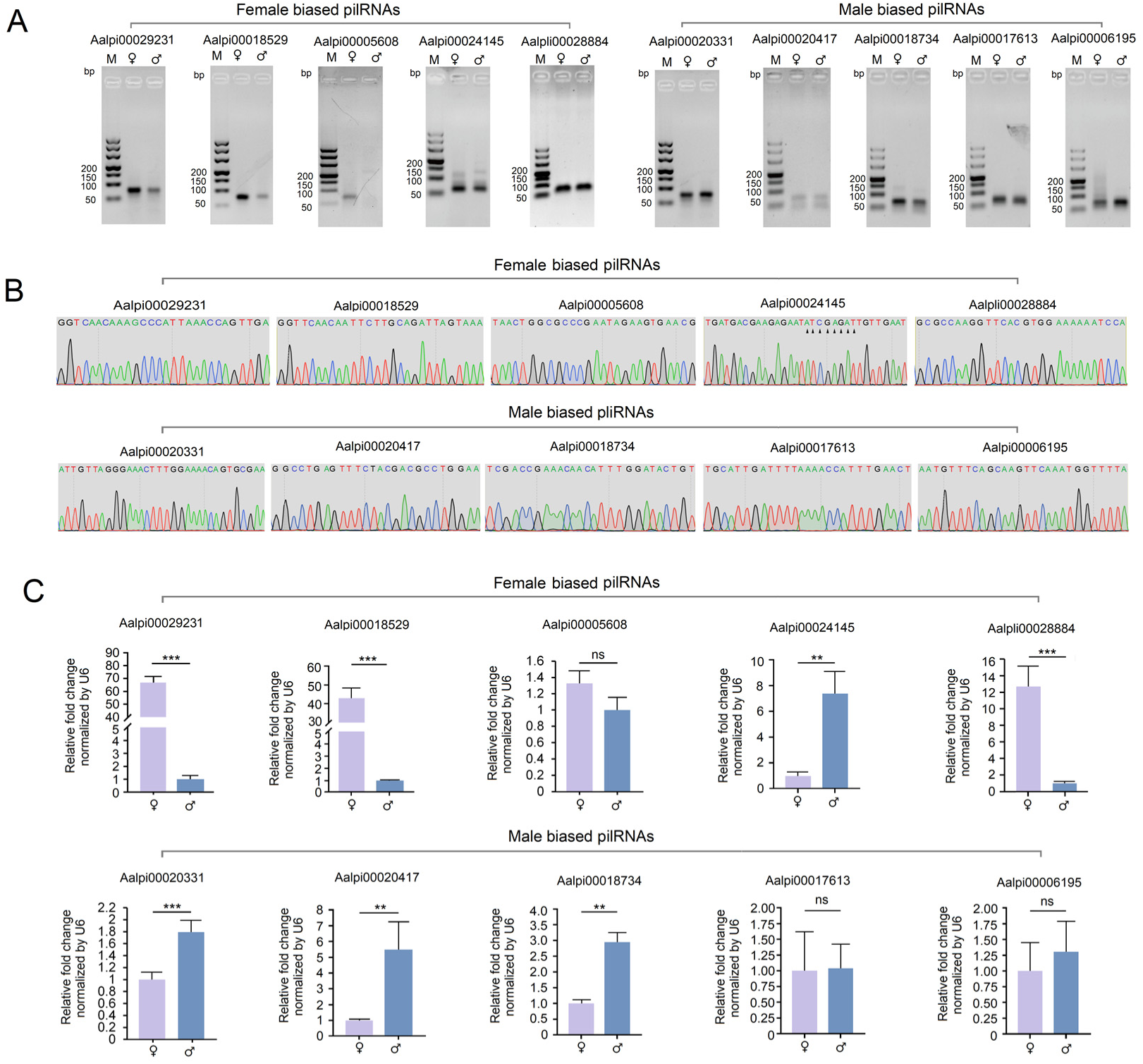
Experimental validation of sex-biased pilRNAs. (A) Detection of ten sex-biased pilRNAs by stem-loop RT-PCR. The PCR products were run on a 1.5% agarose gel in 1X TBE stained with ethidium bromide. (B) Sanger sequencing of PCR products amplified with specific stem-loop primer-based RT-PCR. The triangle indicates eight consecutive base variations detected at nucleotide positions 17-24 of Aalpi00024145. (C) Quantification of ten sex-biased pilRNAs by stem‒loop qRT-PCR. qRT‒PCR was performed in triplicate with three biological replicates. All data are presented as the means ± SEMs, and one-way ANOVA was used to compare the means among different groups. * *P* < 0.05; ** *P* < 0.01; *** *P* <0.001; ns, no significance.

### 3.3 Aalpi-18529 is highly expressed in the ovary and correlates with blood meal

Mature Aalpi00018529 (abbreviations Aalpi18529) is 28 nucleotides (nt) long, with uracil at the 5’ end (1U). Aalpi18529 originates from piRNA Cluster 215 (NW_021837156.1, 62116929-62144956) in scaffold_11 (NW_021837156.1) and is located in exon 15 of the antisense transcript of a lncRNA (TCONS_00030668, XLOC_018874) (NW_021837156.1, 62137804-62137831) (Fig. 2A) [19,30]. Considering that Aalpi18529 has only one transcript without alternative isoforms, we decided to focus our attention on the potential biological functions of Aalpi18529 in female adults.

**Figure 2:**
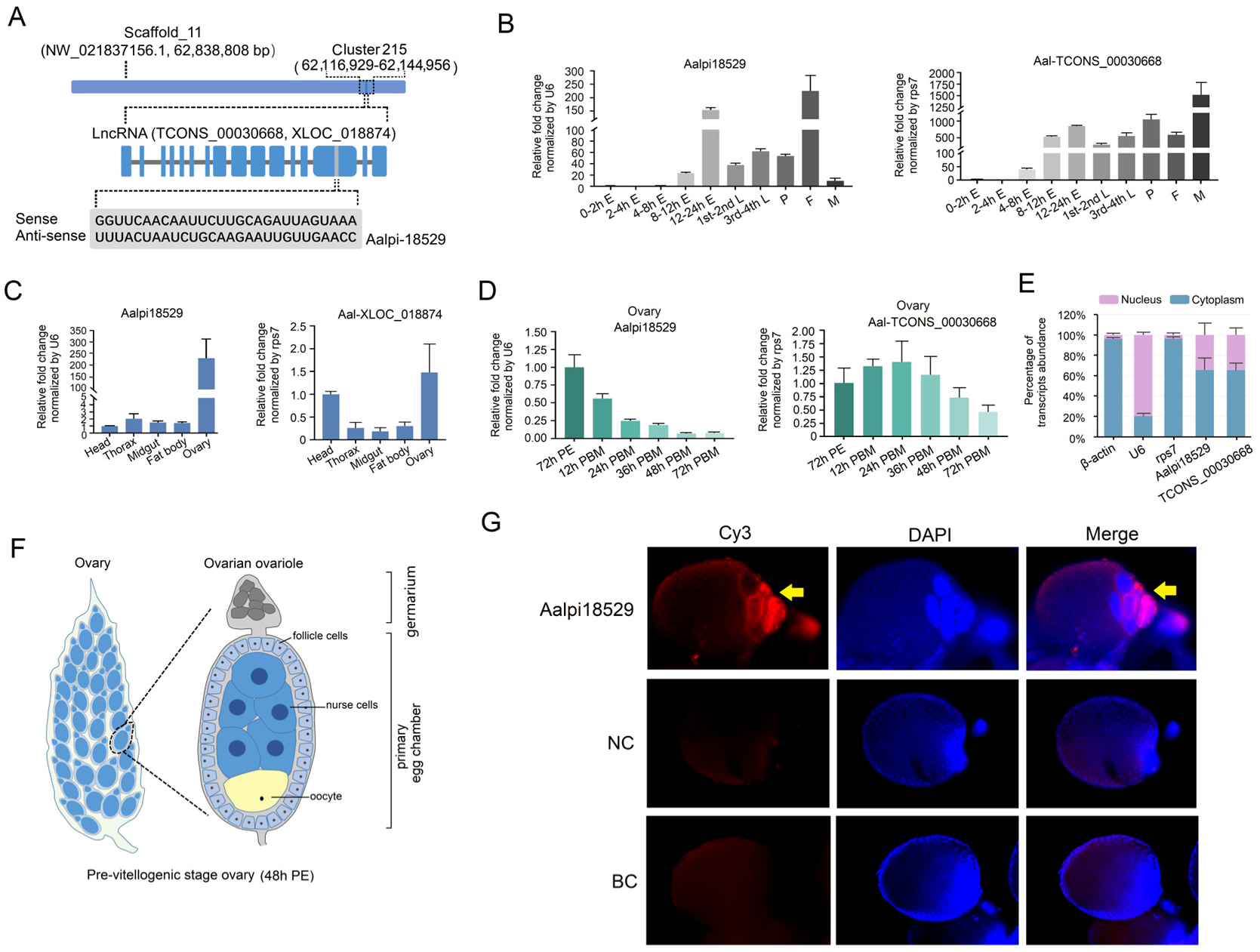
Spatial-temporal patterns of Aalpi-18529 and its host gene measured by qRT‒PCR. (A) Schematic diagram showing the relative position of mature Aalpi00018529 in the *Ae. albopictus* genome. Aalpi18529 originates from piRNA Cluster 215 (NW_021837156.1, 62116929-62144956) in scaffold_11 (NW_021837156.1, length 62,838,808 bp) and is located in exon 15 of the antisense transcript of a lncRNA (TCONS_00030668). (B) Temporal profiles of Aalpi-18529 and TCONS_00030668 at different developmental stages of *Aedes albopictus* determined by qRT‒PCR. E = n hours postoviposition embryo; 1^st^ -2^nd^ L= 1^st^ -2^nd^ instar larvae; 3^rd^-4^th^ L = 3^rd^-4^th^ instar larvae; P = pupae; F = adult female; M = adult male. The relative expression levels of Aalpi-18529 and TCONS_00030668 in 0-2 h postoviposition embryos were set as 1. (C) Spatial expression patterns of Aalpi-18529 and TCONS_00030668 in different tissues of *Ae. albopictus* adult females determined by qRT‒PCR. The relative expression levels of Aalpi-18529 and TCONS_00030668 in the head were set as 1. (D) Dynamic expression levels of Aalpi-18529 and TCONS_00030668 in the ovaries of adult females at different time points after a blood meal determined by qRT‒PCR. PE = hours postemergence; PBM = hours post blood meal. The relative expression levels of Aalpi-18529 and TCONS_00030668 in the 72 h PE groups were set as 1. (E) The relative expression quantities of Aalpi-18529 and TCONS_00030668 in the cell nucleus and cytoplasm fractions of adult females were measured by qRT‒PCR. *AalRps7* and *β-actin* mRNA served as markers of cytoplasmic fractions, whereas *U6* served as a marker of nuclear fractions. The X-axis indicates the different sample groups, and the Y-axis shows the relative expression levels. All qRT-PCRs were performed in triplicate with three biological replicates, and the values are presented as the means ± SEMs. (F) Schematics of the previtellogenic stage ovary and its ovarian ovariole (48 PE), and the major cellular types of the primary egg chamber are indicated. (G) Localization of Aalpi-18529 in the ovaries of adult females. The nucleic acid probe targeting Aalpi-18529 was conjugated to the dual fluorophore Cy3 (red). Probes targeting a scrambled nucleotide sequence were used as the control for piRNA. Egg chambers were stained with DAPI to visualize nurse cell nuclear morphology. BC, blank control; NC, negative control. The images were acquired using an All-in-One Fluorescence microimaging system BZ-X800/BZ-X810 (KEYENCE).

To analyse the temporal expression profiles of Aalpi-18529, total RNA was extracted from various embryonic stages to adults and reverse transcribed with a stem‒loop reverse transcription primer. Subsequently, quantification of Aalpi-18529 was performed by qRT-PCR analysis. The expression pattern of the Aalpi-18529 host gene lncRNA TCONS_00030668 was also determined. Briefly, Aalpi18529 showed the highest relative expression level in the adult female stages and was submaximal in embryos 12-24 h after egg laying, whereas TCONS_00030668 reached its maximum in adult males.

The spatial expression pattern of Aalpi-18529 and TCONS_00030668 in *Ae. albopictus* females was also quantified by performing qRT‒PCR on total RNA extracted from different tissues, including the head, thorax, midgut, ovaries and fat bodies dissected from adult mosquitoes 72 h postemergence (PE). As shown in Fig. 2B, Aalpi-18529 showed relatively obvious ovary-enriched expression patterns and was over 100 times that of other tissues, whereas TCONS_00030668 was predominantly expressed in the ovary and head, which was only a few times that of the other tissues.

To explore whether ovary-enriched Aalpi-18529 was involved in various physiological events in mosquito ovarian development post blood meal, and therefore 72 h PE female mosquitoes were provided a blood meal, ovaries were dissected and collected at different time points post blood meal, and the dynamic expression of Aalpi-18529 during ovarian development was detected using qRT-PCR. As a result, in the ovary, the transcriptional level of Aalpi-18529 showed a continuously decreasing expression pattern post blood meal and was reduced over 12 times at 72 h PBM (Fig. 2B, C). We also compared the expression levels of Aalpi-18529 between virgin and mated female mosquitoes both before and after bloodfeeding to exclude the impact of mating behaviour on piRNA transcriptional level (Fig. S7).

We determined the subcellular localization of Aalpi-18529 in the ovary. Briefly, the ovaries of virgin adult females were dissected, and nuclear and cytoplasmic fraction separation was performed as described in the methods. Levels of Aalpi-18529, TCONS_00030668, *β-actin*, *rps7* and *U6 snRNA* in purified nuclear and cytoplasm fractions were measured by qRT-PCR. As a result, 65.57% of Aalpi-18529 and 65.60% of its host mRNA were detected in the cytoplasm fraction, which indicated that both Aalpi-18529 and its host gene are localized predominantly in the cytoplasm of adult female ovarian cells (Fig. 2D).

Mosquito previtellogenic stage egg chambers comprised of an oocyte, nurse cells and enveloping follicle cells. To detect the Aalpi-18529 transcript localization patterns in different cell types of the ovarian chamber, we designed and synthesized a probe complementary to Aalpi-18529 and performed FISH. The nurse cells were distinguished from the single oocyte-fated cells based on the intensity of DAPI staining as previously described [47]. As a result, Aalpi-18529 primarily localized and concentrated in the cytoplasm of nurse cells and the germarium (the germarium contains germline stem cells (gsc)) and two-cell cysts composed of prospective oocytes (po) and prospective nurse cells (pnc) during the arrest phase (48 h PE) (Fig. 2F, G) [48].

### 3.4 PIWI5 directly binds Aalpi-18529

Although many aspects of piRNA biogenesis in Aedes mosquitoes still need to be further revealed, recent studies indicate that two factors could play a significant role in the primary piRNA pathway and following the Ping-Pong amplification loop, namely, piwi4 and the multiprotein complex Ven body (Fig. 3A) [33,42,49]. To investigate the involvement of PIWI4 and different proteins of Ven bodies in Aalpi-18529 biogenesis, the effects of RNAi-mediated knockdown of *Piwi4* and three Ven body genes, *Ago3*, *Piwi5* and *Vasa*, on Aalpi-18529 expression levels were assessed (Fig. 3B). In addition, for comparison, two Argonaute proteins, AGO1 and AGO2, mainly associated with miRNA and siRNA pathways, were also knocked down by RNAi silencing. The knockdown efficiency of the dsRNA was first assessed by qRT‒PCR. The RNAi results showed significantly reduced *Piwi4*, *Ago3*, *Piwi5*, *Vasa*, *Ago1* and *Ago2* expression to 69.99%, 58.34%, and 78.24% and 74.77%, 42.15%, and 66.64% at 4 d post injection, respectively, compared with the dsRNA-RFP control group (Fig. 3B). As a result, *Piwi5* and *Vasa* knockdown resulted in an obviously decreased abundance of Aalpi-18529, while *Ago3* knockdown caused a mild reduction in Aalpi-18529 (Fig. 3C). However, no significant difference was found in the *Piwi4* knockdown group compared to the control. As expected, *Ago2* interference had no effect on the expression level of Aalpi-18529. Intriguingly, *Ago1* interference also decreased Aalpi-18529 transcript levels. Moreover, the selected AGO subfamily and PIWI subfamily gene interference did not cause a significant difference in the host gene lncRNA TCONS_00030668 (Fig. 3D).

**Figure 3.**
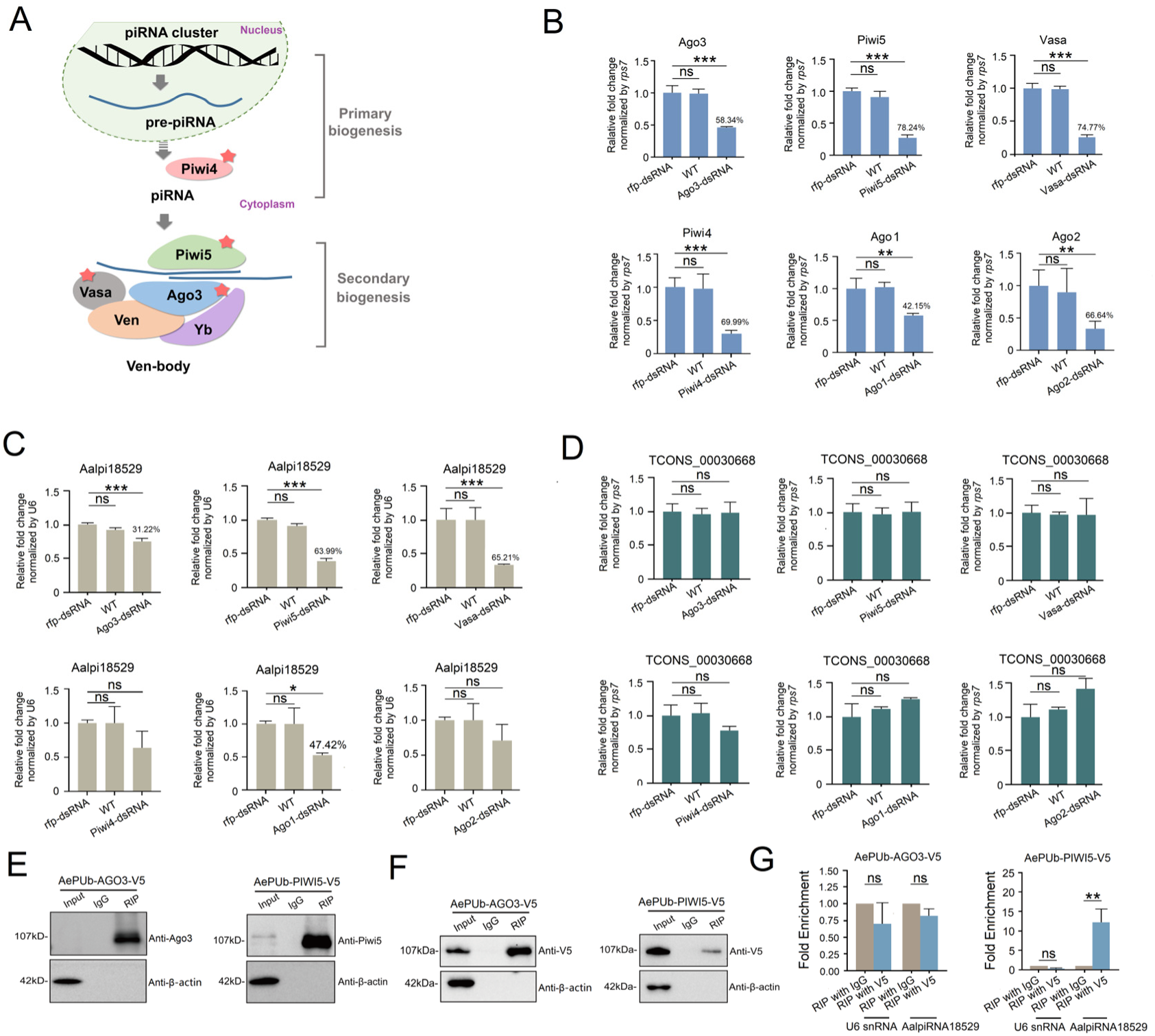
Effect of AGO and PIWI subfamily gene interference on the expression level of Aalpi-18529 and direct interaction between the Aalpi-18529 and AGO/PIWI proteins. (A) Schematic model of the most relevant factors responsible for primary and secondary piRNA pathways in teleosts thus far. The red pentangle indicates the candidate AGO and the PIWI subfamily genes for RNAi. The dotted arrow indicates that the mechanism by which *Piwi4* affects the production of piRNAs still needs further exploration. (B) The knockdown efficiency of *Ago3*, *Piwi5*, *Vasa*, *Piwi4*, *Ago1* and *Ago2* was evaluated by qRT‒PCR. The relative expression levels of Aalpi-18529 (C) and its host gene TCONS_00030668 (D) were determined by qRT‒PCR. (E) Coimmunoprecipitation of PIWI5 and AGO3. The RIP assay was performed in C6/36 cells that express V5-tagged PIWI5 and AGO3 using an anti-V5 tag antibody. Anti-V5 monoclonal antibody and PIWI5/ AGO3 polyclonal were used to pull down the PIWI5/AGO3-V5flag fusion protein, respectively, and then compared and analysed by Western blotting. The lysates before IP were used as a positive control, and IgG pulled down proteins as a negative control. (F) RIP assay was performed using anti-V5 flag antibody with C6/36 cells that express V5-tagged PIWI5 and AGO3. (G) RT‒qPCR assays were conducted to measure the enrichment level of pi18529 in IgG or AGO3/PIWI5 immunoprecipitation complexes. Relative expression in the IgG groups was set as 1. qRT‒PCR was performed in triplicate with three biological replicates. All data are presented as the means ± SEMs, and one-way ANOVA was used to compare the means among different groups. \**P* < 0.05; ** *P* < 0.01; \*\*\**P* <0.001; ns., no significance.

To further determine the direct interaction between Aalpi-18529 and the PIWI protein, we cotransfected the C6/36 cell line with the PIWI5/AGO3 protein fused with the V5-flag and Aalpi-18529 mimics bearing 2ʹ-O-methyl-modified 3ʹ termini. Then, we immunoprecipitated the protein from transfected C6/36 cells using the anti-V5-flag antibody (The specificity of antibodies were confirmed in supplemental figure S6). Subsequently, Western blotting was used to evaluate PIWI5/AGO3 expression and compared the V5-flag monoclonal antibody and PIWI5/AGO3 polyclonal antibody. The WB results revealed that the expressed PIWI5/AGO3-V5flag protein could be recognized by both anti-V5-flag and anti-PIWI5/AGO3 antibodies in RIP samples (Fig. 3E and F), indicating that the V5-flag monoclonal antibody-mediated RIP assay was successfully performed. Levels of Aalpi-18529 in the immunoprecipitates (IPs) were detected by quantitative RT‒PCR. The enrichment of Aalpi-18529 in PIWI5 IPs was increased by more than 10-fold compared with that in the RIP IgG group (Fig. 3G), whereas *U6* showed no significant difference. In contrast, RIP analysis showed that Aalpi-18529 was not enriched in the RIP-AGO3 group (Fig. 3G).

### 3.5 Overexpression of Aalpi-18529 Suppressed Ovarian Development

Considering the distinct spatial expression of Aalpi-18529 in the ovary and its downregulation post blood meal, we further explored the potential function of Aalpi-18529 in the ovary during ovarian development. To overexpress Aalpi-18529 during vitellogenesis, intrathoracic injection with Aalpi-18529 mimics was carried out, and NC mimics were used as a negative control (Fig. 4B). The overall processing workflow is depicted in Fig. 4A. Briefly, 2 days post emergence, virgin adult females were thoracically injected with NC mimics using a nanoinjector. Forty-eight hours post injection, mosquitoes were fed commercial defibrinated sheep blood, and only freshly fed mosquitoes with a fully engorged abdomen with bright red blood were picked out and dissected 72 h PBM per group to analyse the effect of mimics on ovarian development. The overexpression efficiency of the piRNA mimics was assessed by qRT‒PCR. As Fig. 4 shows, the expression of Aalpi-18529 in the ovary was significantly increased by approximately 50 times at 48 h and 72 h post injection, respectively, compared with the NC mimics groups (Fig. 4B). To further confirm the specificity of the overexpression, the expression level of the host lncRNA TCONS_00030668 was also measured by qRT‒PCR. As expected, treatment with piRNA mimics did not influence the transcript level of TCONS_00030668. Then, the piRNA mimic-treated mosquitoes were given a blood meal and dissected at 72 h PBM for phenotypic manifestation observation. As a result, Aalpi-18529 overexpression suppressed ovarian development, and ovarian follicle growth in the piRNA treatment groups was inhibited, with an average primary follicle length of 249.22±11.99 μm being less than that of the WT group (325.96±5.19, n = 45) (*P* < 0.001) and NC mimics group (324.05±5.52, n = 47) (*P* < 0.001). Moreover, the number of developing follicles showed a significant decrease (63.89±4.38, n=47) compared with the WT group (80.51±2.81, n=45) (*P* <0.01) and NC group (79.00±2.26, n = 47) (*P* < 0.001) (Fig. 4C and D).

**Figure 4:**
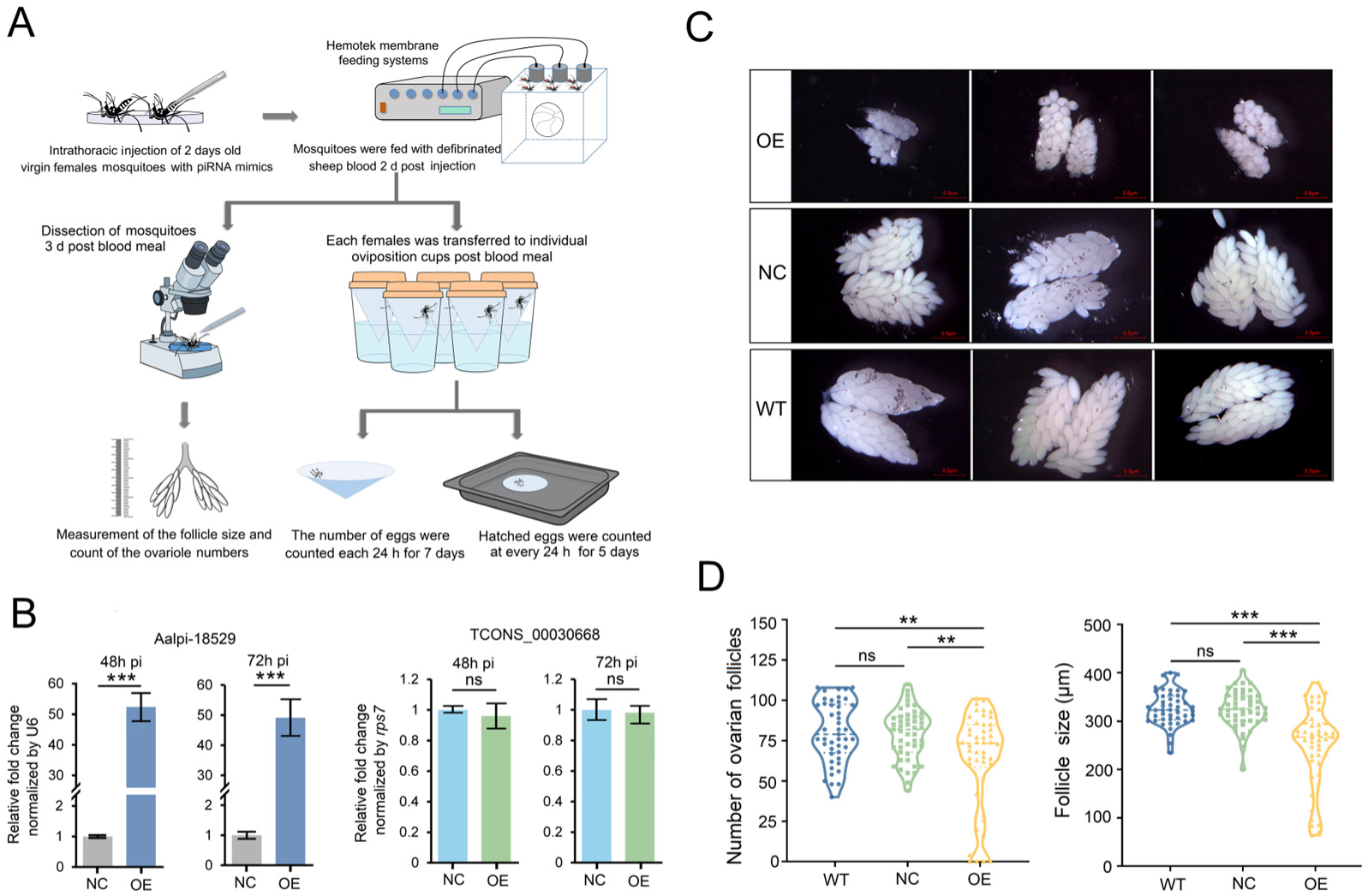
Effect of Aalpi-18529 overexpression on ovarian development in adult female *Ae. albopictus*. (A) Workflow diagram of the general processing analysis of the effect of Aalpi-18529 overexpression on ovarian development, fecundity and egg hatching rate. (B) The relative expression levels of Aalpi-18529 and its host gene TCONS_00030668 were determined by qRT‒PCR. Relative expression in the WT groups was set as 1. qRT‒PCR was performed in triplicate with three biological replicates. (C) Representative images of ovaries dissected from Aalpi-18529 overexpression, NC and WT female mosquitoes at 72 h PBM. All images were taken with a Nikon SMZ1000 stereomicroscope with DIGITAL SIGHT DS-U3 (scale bar: 0.5 μm). (D) Average follicle size (length of the long axis) of ovaries isolated from female mosquitoes and number of developing follicles per female individual in Aalpi-18529 overexpression (n = 46), NC (n = 47) and WT (n = 45) groups. All data are presented as the means ± SEMs, and one-way ANOVA was used to compare the means among different groups. \**P* < 0.05; \*\**P* < 0.01; \*\*\**P*<0.001; ns., no significance.

### 3.6 Overexpression of Aalpi-18529 reduces fecundity and fertility in *Ae. albopictus*

To explore the impact of Aalpi-18529 on female fecundity during the first gonadotrophic cycle, Aalpi-18529 was overexpressed in mated females of *Ae. albopictus* by the intrathoracic injection method. In brief, after copulation, the males were removed, and 2-day-old females were injected with piRNA mimics. Two days post injection, mosquitoes were fed defibrinated sheep blood, and individual oviposition cups were transferred to each mosquito. The number of eggs produced was counted every 24 h for 7 days. The Fecundity assay flow chart is shown in Fig. 4. The overall mean (±SEM) number of eggs oviposited per female at the end of the study period was 48.96 (±3.50), 64.96 (±4.70) and 65.78 (±4.62) for the treatment, control, and WT groups, respectively. As Fig. S3 shows, overexpression of Aalpi-18529 led to a significant decrease of approximately 20% in the average total number of eggs laid by each female compared to the control female (*P* <0.001). As the curves of cumulative egg laying, there was also a significant decrease in the cumulative eggs produced per female adult in the treatment compared to the control at each time point (Fig. S3B, in 4d PBM, 5d PBM and 6d PBM, the *t* values of the paired T test were 2.093, 2.210, 2.229, df = 26, *P* <0.05). In particular, as the main peak period of oviposition (3d PBM), piRNA overexpression caused a nearly 45% reduction compared with the control when comparing the mean number of eggs laid daily (Fig. S3A). Furthermore, total number of eggs laid per group of *Ae. albopictus* adults (n=9 per group) was also decreased compared with control (Fig. S3C). The results indicate that Aalpi-18529 overexpression reduces fecundity in *Ae. albopictus*.

Because piRNAs play an important role in the embryonic development of animals, to investigate whether Aalpi-18529 overexpression has a significant influence on embryogenesis, we also compared the egg hatch rate of piRNA treatments with that of the control. A significant reduction in the egg hatch rate was also observed (Fig. S3D). Moreover, a high level of Aalpi-18529 was detected in the 0-12 h embryo compared to the control (Fig. S3E). These results were consistent with the temporal expression pattern of Aalpi-18529, i.e., Aalpi-18529 maintains relatively low expression levels in early embryonic stages, and overexpressed piRNAs in adult females can serve as maternal RNAs and be deposited into eggs to further influence embryonic development and promote female mosquito fertility decline.

Furthermore, we attempted to determine whether Aalpi-18529 also affects egg phenotypes, including eggshell formation and melanization. The eggs in the treatment and control groups were continuously observed at 0 h, 0.5 h, 2 h, 3 h, 12 h, and 24 h post oviposition (hpo). However, we did not observe any significant phenotypic changes in eggs in response to Aalpi-18529 overexpression (Fig. S3F).

### 3.7 Overexpression of Aalpi-18529 decreases GADD45Α expression in the ovary

To better understand the effect of Aalpi-18529 on ovarian development post blood meal, we performed genome-wide transcriptional profiling analysis of ovaries that were treated with Aalpi-18529 mimics and yielded 240 transcripts that were significantly differentially expressed (|log_2_ fold change| > 0.50 with adjusted *P* value < 0.05); 118 were upregulated and 122 were downregulated in ovaries of Aalpi-18529 mimic-treated females (Table S4). Aalpi-18529 associates with a ping-pong amplification complex, the Ven body, and was found to be mainly enriched in the cytoplasm. We focused on the roles of Aalpi-18529 in posttranscriptional gene regulation. Therefore, we first selected candidate genes involved in pathways related to ovarian development based on KEGG enrichment analyses (Fig. S4) and then measured 14 downregulated transcripts (GenBank ID: LOC115269176, LOC109427191, LOC115261162, LOC109428509, LOC109432989, LOC109430090, LOC115253714, LOC109397705, LOC109415475, LOC109423337, LOC109423833, LOC109432834, LOC109419098 and LOC115266296) by qPCR, and 12 transcripts were found to be consistent with the RNA-seq results. Moreover, as the direct target genes of Aalpi-18529, these genes prefer to have opposite temporal expression patterns compared with piRNAs, so we determined the candidate gene expression levels at various time points after the blood meal, and six genes showed overall increasing expression trends (Fig. 5B). On the other hand, Aalpi-18529 displays a distinct high expression pattern in previtellogenic ovaries, so we injected females with Aalpi-18529 inhibitor and then quantitated the changes in the expression levels of these six genes. As a result, LOC109423337 (Growth arrest and DNA damage-inducible protein, GADD45Α) and LOC115269176 (mitochondrial import inner membrane translocase subunit Tim13-like, TIMM13-like) showed a significant increase compared with the control. Spatial expression pattern of both genes were further analyzed by RT-qPCR, and result revealed that only GADD45a showed enriched expression in the ovary (Fig. S5).

**Figure 5.**
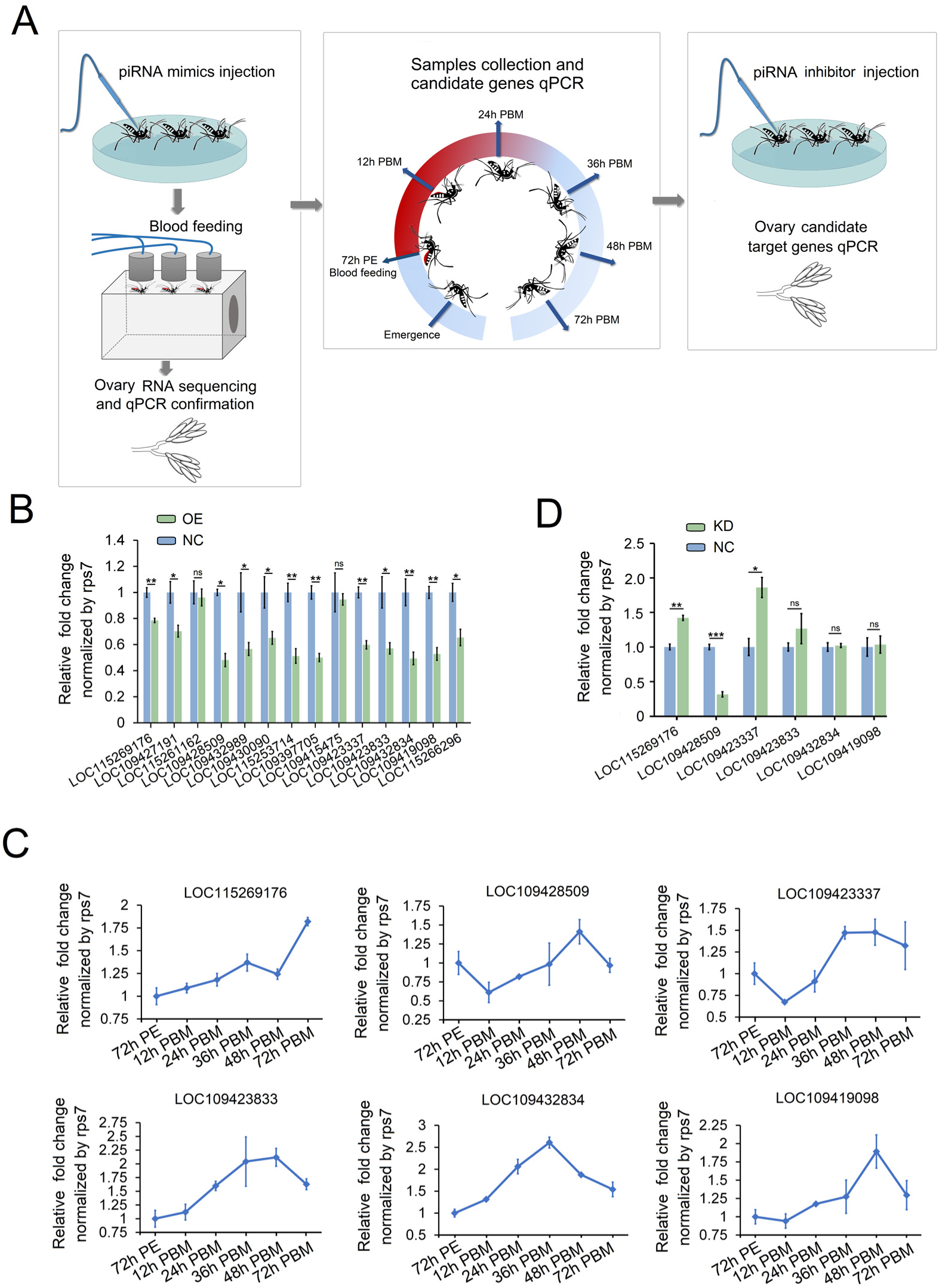
Screen for potential Aalpi-18529 target genes in the ovary of *Aedes albopictus*. (A) Workflow diagram of the general processing analysis of the screen of Aalpi-18529 target genes in the ovary. (B) The relative expression level of identified differentially expressed genes in piRNA overexpression groups by RNA-seq analysis was determined by qRT‒PCR and compared with NC groups. (C) Dynamic expression levels of Aalpi-18529 target genes in ovaries at different time points after a blood meal determined by qRT‒PCR. (D) The relative expressional level of identified differentially expressed genes in piRNA Inhibition groups by RNA-seq analysis was determined by qRT-PCR and compared with NC groups. PE = hours postemergence; PBM = hours post blood meal. The relative expression level of each gene in the 72 h PE groups was set as 1. Different background colours indicate the three phases of the female mosquito oogenesis cycle. All qRT-PCRs were performed in triplicate with three biological replicates, and the data are shown as the means ± SEMs. Student’s *t*-test was used to compare the means between two groups, and one-way ANOVA was used to compare the means among different groups. \**P*<0.05; \*\**P*< 0.01; \*\*\**P*< 0.001; ns, no significance.

### 3.8 *Gadd45a* is the direct target gene of Aalpi-18529

To explore whether Aalpi-18529 regulates ovarian development, fecundity and egg hatching rate by directly targeting *Gadd45a*, RNAhybrid was first used to detect piRNA binding sites (Fig. 6A). As a result, GADD45Α was predicted to harbour a binding site for Aalpi-18529 located in its 3’UTR at position 1038-1066. Therefore, the entire 3’UTR sequence of GADD45Α was cloned and inserted into the pmirGLO plasmid to construct pGLO-GADD45Α-WT, and then the seed sequences for the putative binding site for Aalpi-18529 were mutated to construct pGLO-GADD45Α-MUT (Fig. 6B). Subsequently, pGLO-GADD45Α-WT or pGLO-GADD45Α-MUT was coinfected with Aalpi-18529 mimics or NC mimics into HEK293T cells. As a result, in the group cotransfected with Aalpi-18529 mimics and pGLO-GADD45Α-WT, the relative luciferase activity was significantly repressed compared with that in the NC mimics groups (*p*<0.01; Fig. 6C). When the binding site was mutated, no significant differences were observed in the luciferase activity of cells cotransfected with the piRNA mimics when compared with the NC group (Fig. 6D). These findings suggested a direct interaction between Aalpi-18529 and GADD45Α mRNAs.

**Figure 6:**
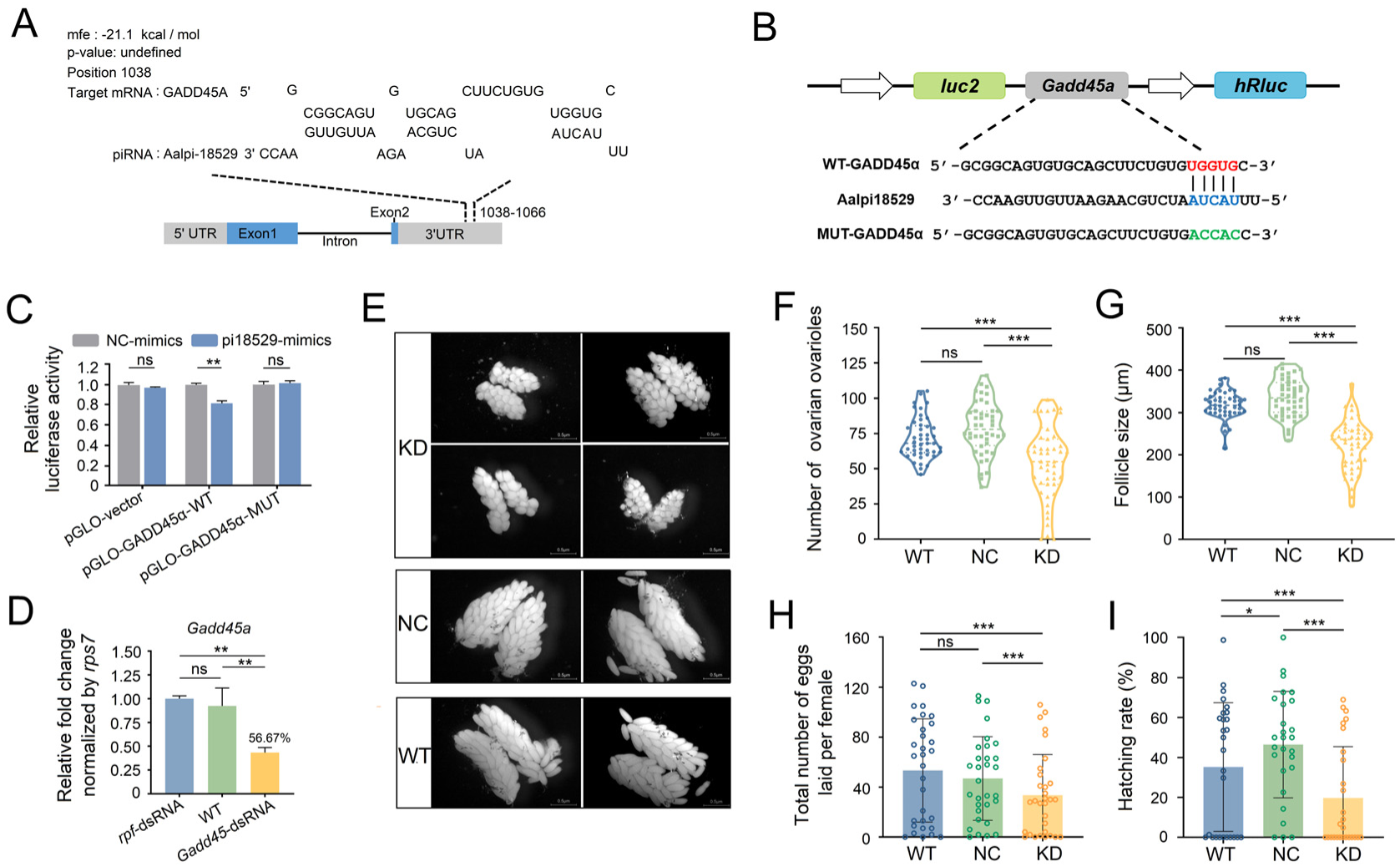
Aalpi-18529 targets the 3’UTR of GADD45A to regulate ovarian and embryonic development. (A) The putative Aalpi-18529 binding site in GADD45Α was predicted by RNAhybrid. (B) Schematic diagram of the dual-luciferase reporter gene structure based on the pmirGLO plasmid. The putative binding sites (WT-GADD45Α) and the mutant version (MUT-GADD45Α) for GADD45Α are presented. (C) Relative luciferase activity was determined 48 h after cotransfecting HEK293T cells with Aalpi-18529/NC (n = 3) and pmirGLO empty vector/GADD45Α-WT/MUT (n = 3). Relative luciferase activity in the NC-mimics groups was set as 1. (D) Relative expression level of *Gadd45a* in female mosquitoes upon thoracic injection of *Gadd45a* dsRNA (KD group), negative control mimics (NC group) and wild-type blank control (WT) determined by qRT‒PCR. qRT‒PCR was performed in triplicate with three biological replicates. (E) Representative images of ovaries dissected from *Gadd45a* KD-treated, NC group and WT group female mosquitoes at 48 h PBM. All images were taken with a Nikon SMZ1000 stereomicroscope with DIGITAL SIGHT DS-U3 (Scale bar: 0.5 μm). (F) and (G) Average follicle size (length of the long axis) of ovaries isolated from female mosquitoes and number of developing follicles per female individual in *Gadd45a* KD (n = 47) NC-mimics (n = 44) and WT (n = 45) mosquitoes. (H) and (I) The total number of eggs laid and hatching rate of eggs laid per *Ae. albopictus* adults. KD (n=27), NC (n = 27) and WT (n = 27). KD, Gadd45a knockdown; NC, negative control; WT, wild type. All data are shown as the mean ± SEM. Student’s *t*-test was used to compare the means between two groups, and one-way ANOVA was used to compare the means among different groups. \*\*\**P* <0.001; ns, no significance.

To investigate whether *Gadd45a* knockdown could phenocopy the phenotype generated by Aalpi-18529 overexpression. DsRNA was designed to target the CDS region of *Gadd45a,* and then 48 h PE female mosquitoes were injected with the dsRNA. The *Gadd45a* knockdown efficiency was validated by qRT‒PCR (Fig. 6E), and adult females were blood fed 48 h post injection. Ovarian development, fecundity and egg hatching rate were assessed according to the method described above. As expected, knockdown of *Gadd45a* analogously resulted in phenotypic defects in ovarian development (Fig. 6F), with a shortened follicle size (Fig. 6G) and a decreased number of developing follicles (Fig. 6H). Moreover, it caused a decrease in fecundity and egg hatching rate (Fig. 6H). Collectively, these results strongly suggest that *Gadd45a* represents a physiological direct target of Aalpi-18529 and plays a significant role in regulating ovarian and embryonic development via *Gadd45a*.

### 3.9 Aalpi-18529 overexpression or *Gadd45a* silencing represses the phosphorylation levels of JNK

GADD45A is a ubiquitously expressed and DNA damage-inducible protein [50,51] that has previously been reported to bind and activate mitogen-activated protein kinase kinase kinase (MAPKKK). MAPKKK activation leads to the phosphorylation and activation of MAPKK, which activates MAPK activity by phosphorylating the threonine and tyrosine residues in MAPK, including c-Jun N-terminal kinases (JNKs) and p38, finally triggering JNK/-dependent apoptosis in mammals [52–54]. A similar role of GADD45Α in JNK-dependent apoptosis was further described in *Drosophila*, which indicated a conserved regulatory cascade in insects [55]. To investigate whether GADD45Α also affects the JNK pathway in Aedes mosquitoes, we first determined the transcript level of *jnk* mRNA in ovaries and embryos treated with Aalpi-18529 mimics and *Gadd45a* dsRNA, respectively. Then, the expression level and phosphorylation level of JNK were detected by Western blotting with a specific antibody. *Jnk* genes of *Ae. albopictus* (GenBank: LOC109429565) encoding three protein isoforms with 334 (GenBank: XP_029719906.1, 38 kDa), 375 (GenBank: XP_029719904.1, 43 kDa) and 377 (GenBank: XP_029719917.1, 43 kDa) amino acids. When using an anti-JNK antibody to detect total JNK or using an anti-phospho JNK antibody to show phosphorylated JNK (pJNK), 43 kDa JNK was the major isoform expressed in each tissue in adult females (Fig. 7F). As Fig. 7 shows, Aalpi-18529 overexpression (Fig. 7B) and *Gadd45a* (Fig. 7C) silencing had no significant effect on JNK transcript levels in ovaries at 48 h, 72 h and 120 h post injection or in embryos (0-24 h post oviposition) at 72 h pi between the treated and control groups (Figure 7D and E). Moreover, no obvious difference in total JNK protein expression levels was observed compared to the control groups (Fig. 7G). However, the phosphorylation of JNK was significantly decreased compared to that in the control groups, as revealed by the anti-phospho JNK antibody (Fig. 7G), we conducted the same experiment in vitro using C6/36 cells, and overexpression of piRNA caused changes in phosphorylation levels of JNK. To further clarify the causal relationship between the decrease in JNK phosphorylation levels and repression of ovary development, we conducted the similar experiment in vitro, and overexpression of Aalpi-18529 also caused the decrease in JNK phosphorylation levels in C6/36 cells (Fig. S8). Which indicating that pJNK was less significantly activated with Aalpi-18529 overexpression and that the level of Aalpi-18529 can influence JNK activation in mosquitoes via the regulation of GADD45Α during vitellogenesis.

**Figure 7.**
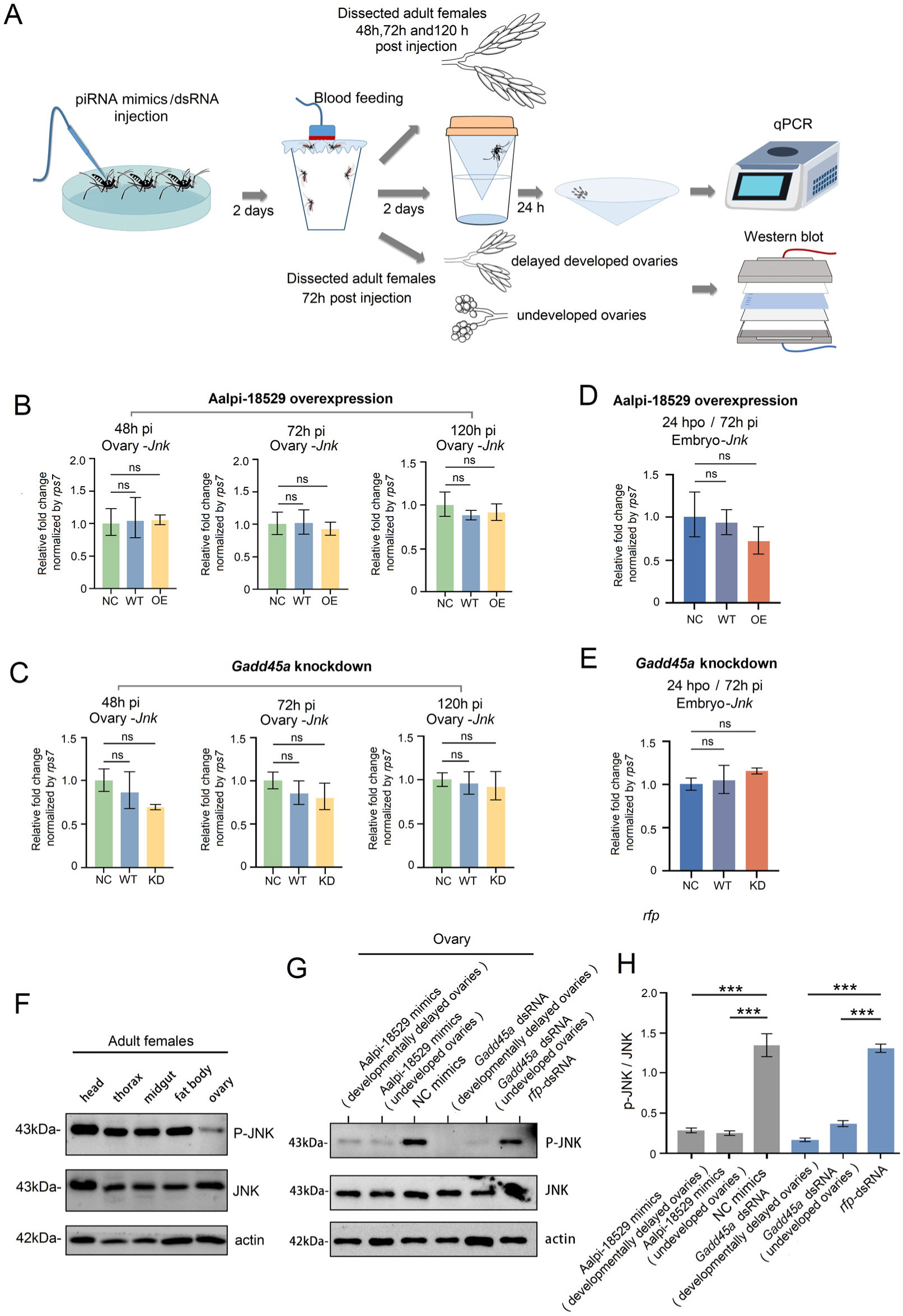
Aalpi-18529 overexpression/*Gadd45a* silencing represses the phosphorylation levels of JNK in *Ae. albopictus*. (A) Workflow diagram of the general processing analysis of the effect of Aalpi-18529 overexpression/*Gadd45a* silencing on JNK phosphorylation. The relative transcriptional level of *ovarian jnk* in the piRNA overexpression groups (B) and *Gadd45a* downregulation groups (C) by RNA-seq analysis was determined by qRT‒ PCR at 48 h, 72 h, and 120 h post injection and compared with the NC and WT groups. The relative transcriptional level of embryonic *jnk* (0-24 h post oviposition) in the piRNA overexpression groups (D) and *Gadd45a* downregulation groups (E) by RNA-seq analysis was determined by qRT‒PCR at 72 h post injection and compared with the NC and WT groups. (F) The spatial expression patterns of pJNK and total JNK in adult females were determined by Western blot using an anti-phospho-JNK antibody and an anti-JNK polyclonal antibody, respectively. (G) Representative immunoblots. Two-day-old female adults were intrathoraxically injected with Aalpi-18529 mimics and *Gadd45a* dsRNA and then fed 2 days later with blood via an artificial blood-feeding system. Subsequently, the ovaries were dissected, and the treatment group ovaries were further divided into two groups, i.e., undeveloped group and delayed developed group. Ovarian proteins were isolated, and protein levels were analysed by Western blot. (H) Semiquantitative analysis of the protein expression of JNK, pJNK and β-actin in the ovary. The density of phosphorylated JNK was quantified with the image analysis software ImageJ and normalized to that of β-actin. *Rps7* was used as an internal control for qPCR, and an antibody against β-actin was used as an internal control in Western blot analyses. triplicates with three biological replicates. All data are presented as the means ± SEMs, and one-way ANOVA was used to compare the means among different groups. \**P* < 0.05; \*\**P* < 0.01; \*\*\**P*<0.001; ns, no significance.

### 3.10 Aalpi-18529 overexpression or *Gadd45a* silencing suppressed nurse cell apoptosis during vitellogenesis

The process of rapid transport of the contents of the nurse cells to the oocyte at the end of vitellogenesis that is accompanied by programmed cell death (PCD, apoptosis) in the nurse cells has been previously described in detail in *D. melanogaster* [56,57], and JNK pathways were thought to be involved in these processes [43,58,59]. To explore whether the decrease in pJNK caused by Aalpi-18529 overexpression leads to the suppression of nurse cell apoptosis during vitellogenesis and ultimately inhibits the ovarian development of *Ae. albopictus,* we performed TUNEL and DAPI staining with ovaries dissected at 24 h PBM. The presence of DNA fragmentation, as depicted by green staining. As a result, Aalpi-18529 overexpression or *Gadd45a* silencing significantly affected ovarian nurse cell PCD in the vitellogenic stage (24 h PBM) (Fig. 8 A) and significantly reduced the number of TUNEL-positive nurse cells in the treatment group by 0.67-fold (Fig. 8B; *p* < 0.05) and 0.50-fold (Fig. 8B; *p* < 0.05) compared to NC mimics and the *rfp*-dsRNA control in the ovary. Furthermore, Aalpi-18529 overexpression or *Gadd45a* silencing also decreased the TUNEL-positive germarium (Fig. 8A). Our results suggest that Aalpi-18529 overexpression alters pJNK-mediated ovarian nurse cell apoptosis during vitellogenesis.

**Figure 8.**
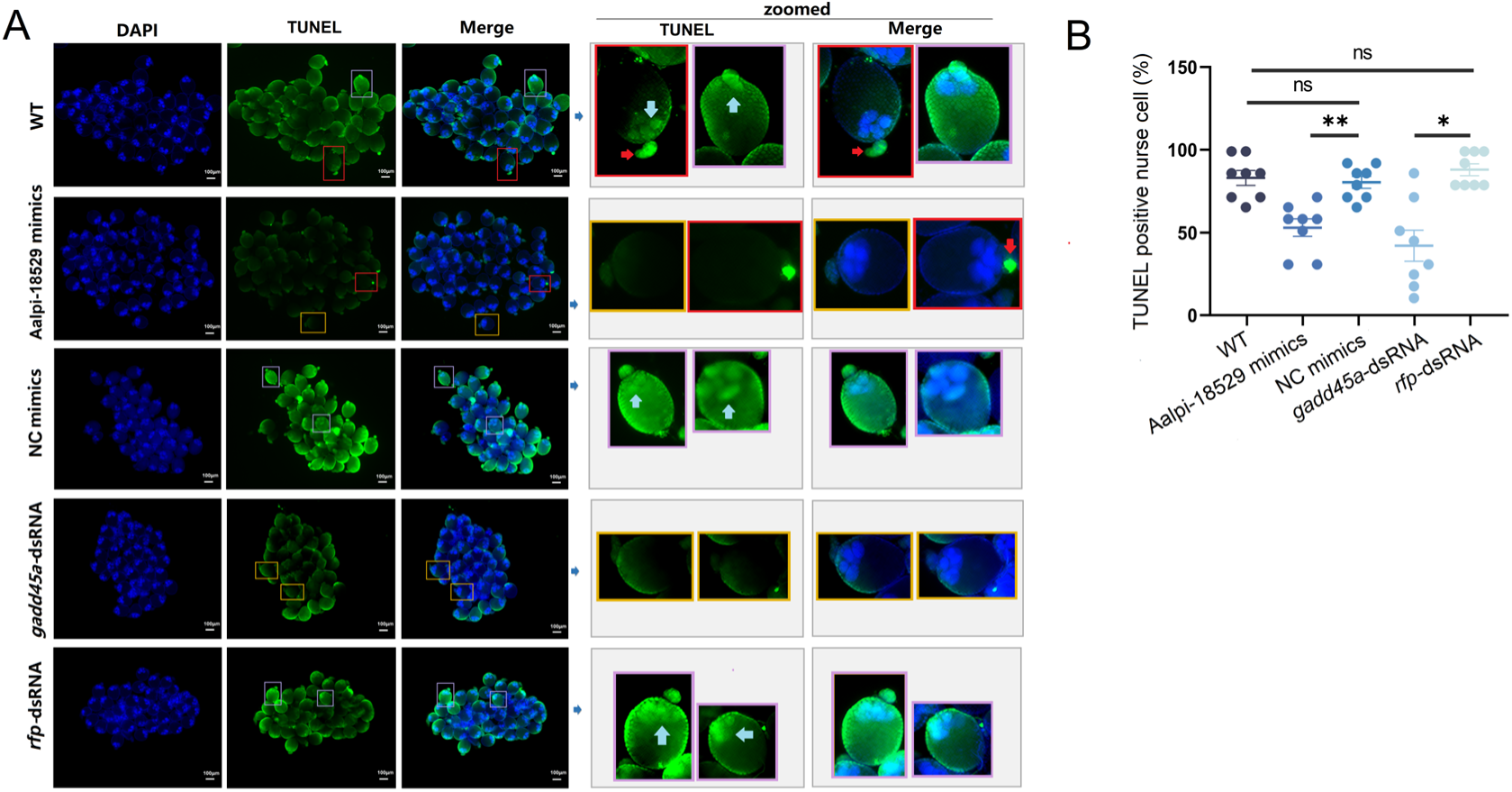
Aalpi-18529 overexpression or *Gadd45a* silencing suppressed ovarian nurse cell apoptosis during vitellogenesis. (A) Ovaries from treatment and control groups were stained with TUNEL assay for apoptotic cells (green), and nuclei were stained with DAPI (blue). Representative pictures of ovaries dissected at 24 h PMB are shown. The right insets show a zoomed image of a typical TUNEL-positive (purpe frame, blue arrow) or TUNEL-negative chamber (yellow frame) and TUNEL-positive germarium (red frame, red arrow). Scale bar: 100 µm. (B) The percentage of TUNEL-positive nurse cells. A total of 15 chambers were randomly selected for TUNEL-positive nurse cell counts in ovaries per adult female, with 8 females per group. All data are shown as the mean ± SEM. Student’s *t-*test was used to compare the means between two groups, and one-way ANOVA was used to compare the means among different groups. \**P* < 0.05. \*\**P* < 0.01. \*\*\**P* <0.001; ns, no significance.

## Discussion

In insects, previous research on the piRNA pathway has been performed mainly in three insects: the fruitfly, silkworm and mosquito. Especially in *Drosophila melanogaster*, the piRNA biogenesis machinery in somatic and germline cells and multifaceted functions in both transposon repression and posttranscriptional regulation have been extensively studied, providing a reference model for piRNA research in other insects [60,61]. More recently, piRNAs research has expanded to locusts, hemipterans, beetles and bees, which has provided broad insights into the potential complexity of the piRNA pathway in the insect kingdom [62]. For mosquitoes, especially culicine mosquitoes, the expansion of the number of piwi genes compared to *D. melanogaster* may be correlated with a larger genome size, greater number of TEs and evolutionary “arms race” between the vector and arbovirus [63]. This finding suggested that piRNA biogenesis in mosquitoes is undoubtedly involved in more complex and intertwined regulatory systems. Hitherto, the exonuclease Nibbler (Nbr) and endonuclease Zucchini (Zuc) were thought to cooperatively determine trailer piRNA production through phased biogenesis, whereas AGO3 and PIWI5, as the core components of the ping-pong amplification loop in Aedes mosquito [64], together with the other factors, Tudor protein Veneno (Ven), VASA and Y-box binding protein (YB), to form a multiprotein complex responsible for ping-pong amplification of piRNAs came from endogenous or exogenous (viral) sequences [42]. In addition, the other Tudor protein Atari that associates with AGO3 has also been confirmed to be involved in an amplification loop. Furthermore, Pasilla protein, as a PIWI5 interactor, is needed for efficient piRNA production [65]. PiRNA-mediated gene silencing was also confirmed in mosquitoes. Target RNA slicing by PIWI proteins may trigger the production of responder and trailer piRNAs through ping-pong amplification and phased piRNA biogenesis, respectively. The piRNAs bind to PIWI5, whereas target gene-derived responder piRNAs load onto AGO3 [66].

Generally, sense-strand piRNAs with an adenine (A) in position 10 (10A bias) tend to bind to AGO3, whereas antisense piRNAs show a strong bias towards a uracil (U) at the 5’ end, preferring to bind PIWI5 [67,68]. Our results show that the mature Aalpi-18529 uniquely matched the antisense lncRNA (XLOC_018874) mRNA and was characterized by 5’ uracil (1U) and was classified as a typical antisense piRNA. By interfering with the major factors AGO3, PIWI5 and VASA, which are involved in ping-pong amplification, the transcriptional level of Aalpi-18529 was significantly depressed. Moreover, the Aalpi-18529 piRNA was detected in PIWI5 IP material, which indicated that Aalpi-18529 biogenesis is tightly associated with PIWI5 and the AGO3-mediated ping-pong cycle, which is in good agreement with the most recent known results for endogenous piRNAs [66].

In *Ae. aegypti*, PIWI4 is also associated with endogenous piRNAs, for instance, satellite repeat-derived tapiR1 [41] and propiR1 detected from Aag2 cells [66]; however, in our work, *Piwi4* interference had no significant impact on Aalpi-18529, which suggests that there are slight variations in piRNA biogenesis. On the other hand, AGO1 knockdown also depressed Aalpi-18529. In mosquitoes, three distinct classes of regulatory sRNA silencing pathways are recognized: siRNA, miRNA and piRNA [69]. In general, miRNAs prefer to interact with AGO1, siRNAs favour AGO2, and piRNAs tend to load onto AGO3. Nevertheless, competition for substrates has been demonstrated previously [70–72]. In *Ae. aegypti Piwi4* was also demonstrated to interact with members of both the siRNA and piRNA pathways [73], highlighting crosstalk among sRNA-mediated RNAi pathways.

Sex-biased gene expression can have a major influence on sexually dimorphic morphological traits, and small RNAs, such as piRNAs, play a large role in controlling gene expression. The piRNA expression profile in mosquito has also made some progress, mainly focusing on the interactions between pathogens and vectors, but research on the differential expression profile of piRNAs with sex bias is relatively limited, especially further functional analysis, lacking basic work [62,74,75]. Our results show that the number of sex-biased piRNAs is enormous, accounting for 30% of the total piRNAs, and these piRNAs play an important role in sexually dimorphic morphological traits in mosquitoes. Aalpi-18529 displays an obvious ovary-enriched expression pattern and is mainly located in the cytoplasm of nurse cells but sharply decreases during vitellogenesis and remains lowly abundant in early embryos, which implies that Aalpi-18529 is not involved in maternal transmission of piRNAs, which can target transposons in early embryos. Overexpression of Aalpi-18529 significantly depressed ovarian development, which also supports our hypothesis that downregulation of Aalpi-18529 in the vitellogenic stage is clearly essential for chamber maturation.

GADD45A is a member of a highly conserved GADD45 family. It is a small acidic protein localized in the nucleus and cytoplasm, and the functional roles of GADD45 have been well elucidated in mammals. GADD45A is involved in various cellular processes, including cell cycle regulation, growth arrest, apoptosis, senescence, and the maintenance of genomic integrity [76,77]. In the apoptosis regulation process, GADD45A first physically interacts with MAPKKK and results in the activation of MAPKKK, which leads to the phosphorylation and activation of MAPKK. Activated MAPKK is thought to further activate its downstream targets MAPKs by phosphorylating threonine and tyrosine residues, including extracellular signal-regulated kinases 1 and 2 (ERK1/2), p38, ERK3/4, ERK5, and JNKs. The mammal’s GADD45 homologue D-GADD45 was also identified in *Drosophila melanogaster* [55]. D-GADD45 plays essential roles in multiple biological processes via the regulation of JNK-dependent apoptosis, for example, affecting egg asymmetric development [55] and proper regeneration of wing imaginal discs [78]. Lifespan [79], and thermal and genotoxic stress resistance [78], which indicated that the GADD45Α-JNK signalling pathway is relatively conserved in animals. Our RNAi and dual-luciferase reporter assay results suggest that GADD45Α of *Ae. albopictus* is the direct target gene of Aalpi-18529, and GADD45Α interference causes a similar phenotype of depression of ovary development. Furthermore, both Aalpi-18529 overexpression and GADD45Α silencing suppressed the pJNK protein level, which suggests that the pJNK decrease in nurse cells during vitellogenesis was the downstream factor in signalling cascades.

In *Drosophila*, D-GADD45 overexpression in the germline affects the dorsal–ventral polarity of the oocyte and causes morphological changes in the dorsal appendages of the eggshell [55]. Although mosquito eggs lack the appendage structures, we also checked the eggshell phenotype of Aalpi-18529 overexpression-and GADD45Α silencing-treated eggs, but we failed to detect any significant phenotypic differences, which suggests that Aalpi-18529 overexpression slows ovarian development but does not affect the final formation of the eggshell of *Ae. albopictus*.

In mosquito ovaries, during the previtellogenic and vitellogenic arrest phase, nurse cells play a key role in oocyte and early embryonic development by delivering transcripts and proteins into the transcriptionally inactive oocyte [80,81]. After a blood meal, the primary egg chambers complete oogenesis by packaging yolk from the fat body as well as RNA and proteins from nurse cells. Meanwhile, nurse cells and follicles begin to undergo developmentally induced apoptosis, and follicle cells secrete eggshell material to give rise to mature egg formation. After oviposition, the secondary egg chambers then become primary egg chambers that enter previtellogenic arrest. Loss of function of Dcp-1 caspase involved in apoptosis results in defects in nurse cell dumping in the *Drosophila* ovary, suggesting that apoptosis is a necessary event for the final transport of the nurse cell contents into the oocyte [56]. Our results showed that Aalpi-18529 is involved in the regulation of apoptosis of nurse cells during vitellogenesis via the GADD45Α/pJNK axis and ultimately affects ovarian development.

In summary, our results showed that an ovary-enriched endogenous piRNA, Aalpi-18529, regulated ovarian development by affecting nurse cell apoptosis in female *Ae. albopictus* after a blood meal via the GADD45Α/pJNK axis. Our study is the first to report a protein-coding gene regulated piRNA by posttranscriptional gene silencing in mosquitoes, expanding our current understanding of the important and multiple roles of piRNAs in biological processes in *Ae. albopictus*. Uncovering the biological functions of sex-biased piRNAs in this species will enhance our understanding of an essential role of the piRNA pathway in mosquito SD and will provide us with more information about the high reproductive capacity of *Ae. albopictus*, which is essential for us to find alternative control strategies for halting dramatic global expansion of this species.

## Acknowledgments

We thank Prof. Zhijian Tu (Department of Biochemistry, Virginia Tech, USA) and Prof. Zhen Zou (Institute of Zoology, Chinese Academy of Sciences, China) for their precious and sincere advices in this study.

## Author contributions

L.Y., Y.H.G., and J.B.G. designed research; L.Y., Y.H.G., Y.L.C., S.Y.R, Y.F.G., and P.W.L. performed research; L.Y., Y.H.G., and Y.L.C. analyzed data; L.Y., K.B. and J.B.G. wrote the paper.

